# Two novel plant-derived yeast species in *Thailandicola* (*Microbotryomycetes*): comparative genomics and transformation

**DOI:** 10.64898/2026.11.19.767971

**Authors:** Julian A. Liber, Marco A. Coelho, Sheng-Yang He

## Abstract

Plant tissues and surfaces are among the largest microbial habitats on Earth, and commensal yeasts are common members of these communities, where they can contribute to plant-microbe interactions including the biological control of plant diseases. Here, we describe two novel species of unpigmented, basidiomycete yeasts, in the class Microbotryomycetes from the plant phyllosphere (*Thailandicola erigeronia* and *T. cardamina*) represented by three isolates from leaves of multiple hosts. We characterize these taxa through analyses of metabolic requirements, tolerance to differences in osmolarity, pH, and temperature, and enzymatic activities. In parallel, we generate near-chromosome-scale hybrid genomes annotated with transcriptome data for these novel species and a congenic isolate. We employ whole-genome and multilocus phylogenetic approaches to infer the placement of these species within a monophyletic clade. We use comparative genomics to examine how the gene content of these yeasts differs from that of other members of the Microbotryomycetes, including an apparent proliferation of retrotransposons. We further demonstrate the genetic transformability of these taxa using *Agrobacterium tumefaciens*-mediated transformation. The description of these new species, together with high-quality genome resources and a genetic transformation protocol, establishes a foundation for experimental studies of these novel plant-derived yeasts and their interactions with hosts and other microbes.

## Introduction

Plant leaves and roots harbor diverse yeast communities that are increasingly recognized as important components of the plant microbiome (Glushakova and Chernov 2010; Boekhout et al. 2022; Sepúlveda et al. 2023). Among these are basidiomycetous yeasts within the class Microbotryomycetes (Pucciniomycotina) including *Microbotryum, Rhodotorula, Sporobolomyces, Rhodosporidiobolus, Leucosporidium, Mastigobasidium, Meredithblackwellia*, *Oberwinklerozyma*, *Microbotryozyma*, *Chrysozyma*, *Colacogloea*, *Heitmania, Slooffia*, *Pseudohyphozyma*, *Fellozyma*, and *Curvibasidium* (Li et al. 2020). Despite growing interest in the application of these yeasts for agricultural (Li et al. 2016) and industrial (Coradetti et al. 2018) uses, continued isolation of previously undescribed taxa indicates that their diversity and biology remain far from fully resolved (Jiang et al. 2024).

Microbotryomycete yeasts are also of ecological and applied interest. This group includes both plant pathogens, such as *Microbotryum* spp. (Hood et al. 2010), and species used for biocontrol of plant pathogens, such as *Rhodotorula* spp. (Castoria et al. 2005). Some members have been investigated for biotechnological applications, including lipid and pigment production (Hernández-Almanza et al. 2014; Coradetti et al. 2018), and genetic tools have been developed for several taxa (Takahashi et al. 2014; Pi et al. 2018; Jiao et al. 2019). They can be abundant in the phylloplane of crops, such as cereals, lettuce, and apples (Last 1955; Hertz et al. 2016; Haelewaters et al. 2021; Kithan-Lundquist et al. 2026).

While the Sporidiobolales (*Rhodotorula*, *Rhodosporidiobolus*, and *Sporobolomyces*) are distinctively pigmented, most other taxa within the class are not readily distinguished by pigmentation. Furthermore, while morphology is more useful in classification of taxon where the teleomorph is observed (Schoutteten et al. 2023), the lack of observed teleomorphs restricts the utility of morphology in distinguishing relatively simple, asexual forms. Earlier classifications relied in part on traits such as cell wall composition and coenzyme Q (CoQ) systems (C. Kurtzman et al. 2011). Beginning with ribosomal sequences (Fell et al. 2000), molecular phylogenetic approaches, often integrated with physiological and metabolic data (Wang et al. 2015; Li et al. 2020; Jiang et al. 2024), are now central to species and genus delimitation in this group. At the same time, expanding genomic resources from initiatives such as the Joint Genome Institute’s Fungal Genomics Program (Grigoriev et al. 2011) have improved phylogenetic resolution (Li et al. 2021) and enabled comparisons of gene content and function relevant to understanding ecological interactions (Martino et al. 2018).

Genetic transformation in basidiomycete yeasts has been pursued with four main approaches: *Agrobacterium tumefaciens*-mediated transformation (ATMT) (Lin et al. 2014), electroporation (Takahashi et al. 2014; Pi et al. 2018), protoplasting (Tully and Gilbert 1985), and gene gun/biolistic (Abbott et al. 2013) methods. Among these, ATMT in fungi, first described by Bundock et al. (1995), has been successfully applied across a broad range of fungi, including ascomycetes (Michielse et al. 2005; Petrucco et al. 2024), mucoromycetes (Ando et al. 2009), chytrids (Medina et al. 2020), and basidiomycetes (Lin et al. 2014). Unlike homology-based methods, the same construct can often be used across multiple strains and/or species, facilitating rapid protocol development. Furthermore, cells do not need to be made competent in the same manner as for electroporation (Pi et al. 2018), which further simplifies the procedure. T-DNA insertion is random (Tzfira et al. 2004) and construct expression may thus vary substantially depending on genomic context.

Here, we propose two novel species isolated from leaves, supported by molecular phylogenetic and metabolic analyses. We also generate near-chromosome-scale genome assemblies, annotated with transcriptome data, and demonstrate successful genetic transformation and heterologous gene expression via ATMT in one of the novel species and one congenic isolate. These near-chromosome-scale assemblies clarify mating systems in this group by suggesting a tetrapolar system with loci on separate telomere-to-telomere contigs.

## Methods

### Sampling

In attempting to isolate a diversity of phylloplane yeasts, we undertook occasional and haphazard sampling of leaf phyllosphere habitats in Durham, North Carolina, USA (N 36.00, W 78.90). We targeted herbaceous plants, including *Erigeron annuus* and *Cardamine hirsuta*, which are common plants in the area. Leaves were collected with sterile forceps into sterile 15 mL conical tubes, which were then processed the same day. Isolates from *Sorghum* roots were obtained as part of Chou et al. (2023).

### Isolation and culturing

Yeast strains were isolated by the ballistospore fall method (Nakase & Takashima, 1993) or by plating ground leaf tissue. For ballistospore fall, a leaf was taped adaxial side-up onto the lid of a 100 mm 1X potato dextrose agar plate (PDA: 39 g/L potato dextrose agar, Difco, supplemented with 34 μg/mL chloramphenicol and 100 μg/mL ampicillin), and placing the agar-containing base over the leaf. The plates were kept in this orientation, sealed with Parafilm, and incubated at room temperature for 3 days, before subculturing on PDA with antibiotics. While forcibly discharged spores (ballistospores) will be captured this way, the short distance between plate and leaf means that electrostatic or other forces could cause yeast propagules to transfer to the agar surface. Yeasts were additionally isolated by harvesting whole leaves, and grinding 1 leaf in 1 mL of 10 mM MgCl_2_ for 3 minutes at 30 Hz in a TissueLyser II (Qiagen) using three ZiO 3 mm beads, then plating 100 μL of undiluted leaf suspension, 10-fold, and 100-fold diluted on PDA1/2 +YE (19.5 g/L Difco potato dextrose agar, 7.5 g/L Bacto agar, and 1 g/L yeast extract, with 34 mg/L chloramphenicol and 100 mg/L ampicillin). Plates were incubated at room temperature and colonies were picked once they appeared, usually 2-3 days. *T. lipolytica* NB124-2 was obtained from the surface-sterilized roots of hybrid *Sorghum*, isolated at Michigan State University and kindly shared with us. These had been isolated by surface cleaning with 0.05% Tween 20, sterilization with 3% hydrogen peroxide, rinses in sterile water, and plating of fine roots into MEA+YE with chloramphenicol, streptomycin, and rifampicin (Chou et al. 2023). Freezer stocks in 50% glycerol were prepared for yeast strains upon identification by ITS sequencing, and maintained at -80°C for long-term storage. Subsequent culturing of strains was performed on YPD (10 g/L yeast extract, 20 g/L peptone, 20 g/L D-glucose, with 15 g/L Bacto agar added for solid media), YM (3 g/L yeast extract, 3 g/L malt extract, 5 g/L peptone, 10 g/L D-glucose, 15-20 g/L Bacto agar), or MEA+YE (10 g/L malt extract, 1 g/L yeast extract, 15 g/L Bacto agar).

### PCR and ITS-LSU sequencing

Polymerase Chain Reaction of the internal transcribed spacer (ITS) region of the rRNA operon, including ITS1-5.8S-ITS2 and partial large subunit (LSU), was performed with Green GoTaq Master Mix (Promega cat. no. M712), using 333 nM of the primers ITS1f (5’-CTTGGTCATTTAGAGGAAGTAA-3’;Gardes & Bruns, 1993) and LR3 (5’-CCGTGTTTCAAGACGGG-3’;Vilgalys & Hester, 1990), with yeast cells added directly to 12 μL reactions with a pipette tip. Cycling parameters were: 95°C for 10 min, 35 cycles of 95°C/30 s, 57°C/30 s, 72°C/1 min, followed by a 5 min hold at 72°C. Amplification of additional LSU sequence was performed with LR0R (5’-ACCCGCTGAACTTAAGC-3’) and LR7 (5’-TACTACCACCAAGATCT-3’), using 54.3°C as the annealing temperature. PCR products were visualized via gel electrophoresis on a 0.9% agarose gel, run for 12 minutes at 150 V and stained with SYBR Safe. PCR products were cleaned of primers and dNTPs with 0.24 U thermolabile exonuclease I (NEB cat. no. M0568) and 0.024 U QuickCIP (NEB cat. no. M0525) in 0.042x rCutSmart buffer and 0.043x NEBuffer r3.1 buffer, with 1.5 μL of PCR product in a 2.7 μL reaction. This mix was heated to 37°C for 15 minutes, then to 80°C for 1 minute. Cleaned PCR products were sequenced by Eurofins Genomics using the ITS1f primer for the ITS region and both LR0R and LR7 for the LSU region. ITS sequences (spanning ITS1f-LR3, including partial SSU, ITS1-5.8S-ITS2, and partial LSU) were searched against GenBank nucleotide and GlobalFungi (Větrovský et al. 2020) databases to attempt identification and determine the frequency of detection or isolation of related strains.

### Genome and transcriptome sequencing

Genomic DNA for whole-genome sequencing was extracted using the Wizard Genomic Purification Kit (Promega cat. no. A1120), modified by omitting enzymatic cell wall degradation and instead lysing cells at 65°C for 1 h in Nuclei Lysis Solution. For isolates *T. erigeronia* JL201 and *T. cardamina* JL221, 650 Mbp of short-read sequencing was generated by SeqCenter (Pittsburgh, PA, USA) on the Illumina NextSeq 2000 platform using paired-end 2 × 151 bp chemistry. Additional long-read sequencing was performed for all three species using Oxford Nanopore Technologies (ONT) v14 library preparation and R10.4.1 flow cells with an expected output of 360 Mb of sequence generated for each by Plasmidsaurus (Eugene, OR, USA). *T. lipolytica* NB124-2 was further sequenced by Plasmidsaurus on the Illumina NextSeq 2000 platform with 2 × 151 bp chemistry, producing 113.6 Mbp of output. A transformed line of *T. erigeronia* JL201 containing a targeted genomic insertion was also sequenced using ONT; reads aligning to the inserted sequence with Minimap2 were removed, then pooled with reads from the wild-type strain prior to assembly.

RNA-Seq was performed on *T. erigeronia* JL201 to examine transcriptional responses to a variety of conditions, including salt stress (5% w/v NaCl), osmotic stress (20% w/v sucrose), pH stress (pH 3.5 or 7), redox stress (1.5 mM H_2_O_2_), nitrogen deprivation (0% ammonium sulfate), and carbon source shift (10 mM citrate and 0 % D-glucose), each compared to a standard synthetic defined medium (SD) containing 1x yeast nitrogen base without amino acids or ammonium sulfate (Millipore-Sigma cat. no. Y1251, 1.7 g/L), 0.5% ammonium sulfate, and 2% D-glucose, pH ∼4.8.

Briefly, a starter culture of 10 mL YPD in a 50 mL conical tube was inoculated from a single colony of *T. erigeronia* JL201 and grown for 2 days at 28°C and shaking at 225 rpm. The culture was diluted to 1050 mL of SD medium and aliquoted in 10 mL volumes into 15 mL conical tubes. These tubes were incubated as before, tilted slightly. After 48 h, the tubes were centrifuged at 3214 × g for 5 min at room temperature, then the T=0 h samples were snap frozen in liquid nitrogen. For the 1 h and 24 h time points, cells were transferred to 2 mL microcentrifuge tubes, then 1.5 mL treatment or control medium was added. After 1 h and 24 h of incubation in the same conditions, the cultures were centrifuged at 15000 × g for 1 min, the supernatant removed, and 450 μL of Qiagen RLT buffer with 1% ꞵ-mercaptoethanol was added before snap freezing. RNA was extracted using a hot acid phenol protocol (Collart and Oliviero 1993), treated with TURBO DNA-free DNase, and sequenced on an Illumina NovaSeq 6000 S4 lane with 2 × 150 bp format following polyA enrichment. The resulting sequencing reads allowed for annotation of gene positions.

### Genome assembly, annotation, and comparative genomic analyses

Genome and transcriptome data are available under BioProject PRJNA892096. Illumina sequencing reads were quality filtered with fastp (Chen et al. 2018). Long reads were assembled with Flye v2.9.2 (Kolmogorov et al. 2019), and the resulting assemblies were then polished with Polypolish v0.6.0 (Bouras et al. 2024) using the Illumina reads aligned with BWA-MEM (Li 2013). Telomeres were annotated with tidk v0.2.0 (Brown et al. 2025). Gene models for JL201 were predicted using BRAKER3 v3.0.6 (Gabriel et al. 2024) implementing AUGUSTUS (Stanke et al. 2006) and GeneMark-ETP (Brůna et al. 2024) on the EU Galaxy platform (Jalili et al. 2020) integrating subsampled mRNA reads following alignment to the genome with HISAT2 v2.1.0 (Kim et al. 2019). The protein set predicted for JL201 was then used as evidence for gene prediction in the remaining genomes. Predicted proteins were functionally annotated with InterProScan (v5.59-91.0), implementing TIGRFAM, FunFam, SFLD, SUPERFAMILY, PANTHER, HAMAP, PROSITE, SMART, PRINTS, PIPSR, PROSITE, AntiFam, MobiDBLite, PIRSF, and UniProtKB.

Comparative genomic analyses based on InterProScan annotations generated for the three *Thailandicola* species and for proteomes of 12 other Microbotryomycetes taxa (<u>Table S1</u>). The relative abundance of annotation terms in *Thailandicola* species compared the other Microbotryomycetes, as well as among *Thailandicola* species, was evaluated using row-wise Fisher’s tests, implemented in rstatix v0.7.2 (Kassambara 2023), with *P* values adjusted with the Holm-Bonferroni method (Holm 1979).

Linear synteny plots of full contigs and mating-type (*MAT*) loci regions were generated with EasyFig v2.2.2 (Sullivan et al. 2011) using BLASTn for alignment. *MAT* loci were first identified by BLAST searches using previously characterized Microbotryomycetes *MAT* genes and their flanking genes as queries (Coelho et al. 2010; Maia et al. 2015). The *HD* loci were defined as the regions spanning the *HD1* and *HD2* genes. The *P/R* loci were preliminary delineated based on structural comparisons between opposite mating types from different species, rather than between opposite mating-type strains of the same species, because additional strains were not available. Accordingly, the inferred boundaries and gene content of the P/R loci should be regarded as provisional.

### Phylogenomic and multilocus phylogenetic analyses

Phylogenetic relationships within the Microbotryomycetes were inferred using a two-step approach in which a genome-scale ASTRAL species tree, derived from whole-genome data, was used to constrain a seven-locus maximum-likelihood phylogeny inferred with IQ-Tree. Whole-genome shotgun assemblies were downloaded from GenBank and JGI Mycocosm, then BUSCO v6.0.0 (Tegenfeldt et al. 2025) was run using the Microbotryomycetes_odb12 profiles on these and the generated genome assemblies to extract nucleotide sequences. MAFFT v7.543 was then used to align nucleotide sequences, followed by gene-tree inference with IQ-Tree v2.2.2.7 (Minh et al. 2020). The species tree was then inferred using weighted ASTRAL v1.23.3.7 (Zhang et al. 2025).

Because most prior studies of the taxonomy of this group (Wang et al. 2015; Li et al. 2020; Jiang et al. 2024) used multi-locus data from rDNA (SSU, LSU, 5.8S) and four protein-coding loci (CYTB, TEF1, RPB1, and RPB2), these loci from type or reference genome strains in the Microbotryomycetes, Agaricostilbomycetes, Spiculogloeomycetes, Mixiomycetes, and Atractiellomycetes were incorporated into the species tree. Sequences from genomes were found using HMMs with HMMER v3.3.2 (Eddy 2011) or ITSx v1.1.3 (Bengtsson-Palme et al. 2013) trained on known sequences. Accessions and genomic coordinates are in <u>Table S2</u>. Each rRNA region and protein-coding locus was aligned using MAFFT, corrected and trimmed manually, and assigned to partitions. Each codon position for each protein-coding gene was assigned to a partition. ITS1 and ITS2 loci as well as intronic sequence from protein-coding loci were excluded from phylogenetic analysis due to poor alignment quality.

ModelFinder (Kalyaanamoorthy et al. 2017) implemented in IQ-Tree v2.2.2.7 (Minh et al. 2020) was used to select the best substitution model for each partition, then tree-inference was performed using the IQ-Tree algorithm followed by 1000 Ultra-Fast bootstraps (Minh et al. 2013). The Agaricostilbomycetes were designated as the outgroup. Trees were manipulated and displayed in FigTree v1.4 (Rambaut 2012) as with ggtree 3.8.2 (Yu et al. 2017). Gene trees for each locus were inferred using the same methodology.

### Yeast Phenotypic Characterization

Colony morphology was examined on YMA (5 g/L peptone, 3 g/L yeast extract, 3 g/L malt extract, 10 g/L D-glucose, 20 g/L agar) after 4 days of incubation at 25°C (Suh et al. 2008). Images were color corrected using a Calibrite color checker card in Adobe Photoshop. Cell morphology was examined by incubating cells in YMB (YMA without agar) for 5 days at 25°C.

Tests for carbon source assimilation and oxidation were performed with Biolog YT Microplates (Biolog cat. no. 1005), using cells grown on BUY agar plates (Biolog cat. no. 71005) for 2 days at 28°C, and resuspended in water to an optical density matching that of the YT standard (Biolog cat. no. 3415). Plates were incubated for 72 h at 28°C, with absorbances measured each day. While assimilation tests can be performed up to 3 weeks (Suh et al. 2008), the microplates used for Biolog assays are recommended by the manufacturer to be observed up to 72 h, likely due to their propensity to dry out or become contaminated with longer incubation. Only carbon assimilation tests were subsequently used, but both assimilation and oxidation tests were assessed by absorbance at 590 nm (for assimilation) and 500 nm (for oxidation). The absorbance for each well was compared to the maximum absorbance for the assimilation tests, with the negative control well set to zero. A substrate scored as positive for assimilation for absorbances ≥50% of the maximum, and was counted as weak for absorbances ≥20% and <50% of the maximum. Substrates were counted as negative if the absorbance was <20% of the maximum (Haelewaters et al. 2020).

All other metabolic and physiological tests, including nitrogen assimilation, vitamin requirement, osmotic and chemical stress tolerance, and enzyme assays were performed according to Suh et al. (2008). Carbon sources were prepared at concentrations of 5 g/L in 6.7 g/L yeast nitrogen base without amino acids (BD cat. no. 291940), 45 mg/L yeast extract, and 45 mg/L casamino acids, and were sterilized using 0.22 μm filters. Soluble starch was autoclaved instead of filtered. ∼100 μL of methanol and 135 μL of absolute ethanol were added to 4.5 mL of pre-filtered base medium for their respective assimilation tests. DL-lactate and citrate were added as 5g/L of their conjugate acids, and their pH was adjusted to 5.0 with NaOH. D-lactose was additionally tested for *T. erigeronia* and *T. cardamina*. A positive control of D-glucose and a negative control without an added carbon source were included. We had previously validated the correlation of the Biolog method and the method of testing in synthetic defined media (Bian et al. 2026). Fermentation substrates were tested at 20 g/L in 10 g/L yeast extract. ∼18 mL of a given medium was added to a 20×150 mm glass test tube, containing an inverted 12×75mm test tube. The inner test tube was manipulated to remove any bubbles, then the tubes were capped and autoclaved for 15 minutes. Tested substrates included D-glucose, D-xylose, D-galactose, D-lactose, and D-sucrose with a negative control of no carbon source. *Saccharomyces cerevisiae* BY4741 was added to a D-glucose test tube to serve as a positive control. Both carbon assimilation and fermentation tests were inoculated by collecting cells of a 2 day-old YMA plate culture, grown at 25°C, suspending cells in water, and adding 100 μL of cell suspension per tube. Tubes were incubated at 25°C without shaking. Turbidity was observed for assimilation tests at 7 and 21 days. Fermentation tests were checked at 7, 14, and 21 to observe bubble formation.

Nitrogen assimilation assays were performed with the following base medium. 2x yeast carbon base was prepared with 20 g/L glucose, 40 μg/L biotin, 4 mg/L calcium pantothenate, 4 μg/L folic acid, 20 mg/L *myo*-inositol, 0.8 mg/L niacin, 0.4 mg/L *para*-aminobenzoic acid (PABA), 0.8 mg/L pyridoxine HCl, 0.4 mg/L riboflavin, 0.8 mg/L thiamine HCl, 2 mg/L L-histidine, 4 mg/L L-methionine, 4 mg/L L-tryptophan, and the following salts, per anhydrous amount: 1 mg/L H_3_BO_3_, 80 μg/L CuSO_4_, 0.2 mg/L KI, 0.4 mg/L FeCl_3_, 0.8 mg/L MnSO_4_, 0.4 mg/L Na_2_MoO_4_, 0.8 mg/L ZnSO_4_, 1.7 g/L KH_2_PO_4,_, 0.3 g/L K_2_HPO_4_, 1 g/L MgSO_4_, 0.2 g/L NaCl, 0.2 g/L CaCl_2_. Nitrogen sources were prepared separately at the following concentrations in 2x yeast carbon base: 0.6 g/L KNO_3_, 0.84 g/L NaNO_2_, 1.33 g/L L-lysine HCl, 1.08 g/L cadaverine HCl, 0.8 g/L creatine monohydrate, 0.68 g/L creatinine, 1.33 g/L D-glucosamine HCl, 0.40 g/L imidazole, and 1.28 g/L D-tryptophan. pH was adjusted to 5.5-6.5 with 3.7% HCl if needed, then 20g/L (final concentration) Bacto agar was added and an equal amount of water to 2x yeast carbon base. The media were then autoclaved and poured into 100 x 15 mm plates.

Vitamin requirement tests were performed using a base medium of 10 g/L D-glucose, 5 g/L Difco vitamin-free casamino acids, 1 g/L KH_2_PO_4_, 0.5 g/L MgSO_4_-7H_2_O, 0.1 g/L CaCl_2_-2H_2_O, and 0.1 g/L NaCl. This base solution was autoclaved, then filter-sterilized vitamin solutions were added at 10x concentration to achieve the same final concentrations as in the 1x nitrogen assimilation medium. These vitamins were biotin, calcium pantothenate, folic acid, *myo*-inositol, niacin, PABA, pyridoxine HCl, riboflavin, and thiamine HCl. Dropout medium without *myo*-inositol, calcium pantothenate, biotin, niacin, pyridoxine HCl, thiamine HCl, or PABA, singularly, and without biotin and thiamine HCl or pyridoxine HCl and thiamine HCl, doubly, were prepared. Cells from a YMA plate grown at 25°C for 2 days were suspended in the base medium for 5 days at 25°C to starve them of vitamins, before adding 100 μL to each culture tube with 5 mL of base medium with all vitamins, dropout medium missing 1 or 2 vitamins, or a negative control without any vitamins. Turbidity was assessed at 3, 7, and 14 days post inoculation.

Tolerance tests were performed on solid medium: acetic acid tolerance (1% glacial acetic acid added to 100 g/L D-glucose, 10 g/L tryptone, 10 g/L yeast extract, 20 g/L Bacto agar after autoclaving and cooling to 60°C), and the following osmolytes in 1% yeast extract and 2% Bacto agar medium: 50% (w/v) D-glucose, 60% (w/v) D-glucose, 10% (w/v) NaCl, 16% (w/v) NaCl, and a control with no osmolyte. A 10-fold dilution series from 1 to 10^6^ dilution of yeast cells grown on YMA plates for 3 days at 25°C and suspended in sterile water was prepared, and 10 μL drops were added to each of the tolerance media. Tolerance tests of high temperatures (30°C and 35°C) and cycloheximide (0.1% and 0.01%, at 25°C) were inoculated using the same cell suspension, and each was done in a solution of 6.7 g/L yeast nitrogen base without amino acids (BD cat. no. 291940) and 5 g/L D-glucose. Growth for tolerance tests was observed at 8 days post inoculation. pH range tolerance was tested using 1.7 g/L yeast nitrogen base without amino acid and ammonium sulfate (Millipore-Sigma cat. no. Y1251) with 5 g/L ammonium sulfate and pH adjusted to 3, 4, 6, 8, or 9 with hydrochloric acid or sodium hydroxide solutions prior to filter sterilization. Growth was assessed in 200 μL volumes in a 96-well microplate, after 4 days of incubation at 28°C.

Urease activity was tested in Urea R Broth (Rustigian and Stuart 1941), composed of: 20 g/L urea, 0.1 g/L yeast extract, 0.01 g/L phenol red, 0.095 g/L Na_2_HPO_4_, 0.091 g/L KH_2_PO_4_, with pH adjusted to 6.9 and filter sterilized. A loop of cells from a YMA plate grown at 25°C for 2 days was added to 0.5 mL of broth, which was then incubated at 37°C for up to 4 h or until color change compared to uninoculated broth was observed. A bright pink or red color indicated urease activity.

Gelatin liquefaction was assessed in a gelatin medium composed of 20% gelatin, 5 g/L D-glucose, and 6.7 g/L yeast nitrogen based. The medium was autoclaved and 2.5 mL was dispensed into culture tubes. Cells from a YMA plate grown at 25°C for 2 days were picked up with a long wood streaker and stabbed into the solidified medium. The culture tubes were then incubated at 25°C for 3 weeks. An uninoculated tube was simultaneously incubated for comparison.

Ballistospore formation was assessed by streaking each culture on a 100 x 15 mm corn meal agar (CMA; Criterion Dehydrated Culture Media, Hardy Diagnostics, Santa Maria, CA, USA) plate, and placing another CMA plate with a sterilized microscope slide below the streaked plate, then sealing the two plates with surgical tape (C.P. Kurtzman et al. 2011). These plates were incubated at room temperature, approximately 22°C, for 3 weeks, and examined for colony formation on the unstreaked plate. Crosses between JL221 and JL257 were attempted on V8, PDA, and YMA media by co-inoculating a light lawn of cells and incubating at room temperature in the dark up to 6 months.

### Genetic Transformation via *Agrobacterium tumefaciens*

Genetic transformation of yeast isolates was performed using an ATMT approach modified from Bundock et al. (1995). *Agrobacterium tumefaciens* AGL-1 was transformed via electroporation (Wolfram Jwd Debler 2017; Wolfram Jwd Debler 2019) with a binary vector plasmid (pMBM51) carrying a aphA1 (NEO) cassette for G418 resistance and enhanced yellow fluorescent protein (eYFP), both driven by the bidirectional histone H3-H4 promoter from *T. cardamina* JL221. This plasmid was originally derived from NM9-SpCas9-NLS3 (Schultz et al. 2019). NM9-SpCas9-NLS3 was a gift from Huimin Zhao (Addgene plasmid # 128177).

Yeasts were grown for 2 days in YPD at 28°C with shaking at 230 rpm in 10 mL culture tubes, then washed with water, diluted to an OD600 = 0.6. Agrobacterium cells were grown for 2 days in LB supplemented with kanamycin (50 mg/L) and rifampicin (20 mg/L) under the same conditions, then washed and diluted to an OD600 = 2.0-6.0. For co-cultivation, 50 μL each of yeast and Agrobacterium cells were added to each well of a 12-well microtiter plate containing 2 mL of 2% agar medium formulated in Bundock et al. (1995) at pH 5.2, with 100 μM acetosyringone, and three 3 mm glass spreading beads. Plates were gently agitated to spread the cells, then allowed to dry. The plates were sealed and incubated at 25°C for 3 days. After incubation, cells were resuspended in 300 μL of sterile water and plated on YPD containing 100 mg/L G418, 50 mg/L tetracycline, and 34 mg/L chloramphenicol. Fluorescent colonies were picked after 3-5 days of incubation at 28°C. T-DNA insertion locations were mapped using the splinkerette PCR method (Potter and Luo 2010) using BstYI digestion and confirmed by PCR across the insertion region.

## Results

### Yeast Isolation and preliminary identification

Three isolates (JL201, JL221, and JL257) with similar colony morphology were independently isolated from the leaves of *Erigeron annuus* (JL201) and *Cardamine hirsuta* (JL221 and JL257) in Durham, NC, USA on August 27, 2021, October 27, 2021, and January 25, 2022, respectively. Isolates JL221 and JL257 were isolated at nearby locations approximately 260 m apart. JL201 was co-isolated with yeasts in *Sporobolomyces*, JL221 with an isolate in *Aureobasidium*, and JL257 with *Hannaella*, *Sporobolomyces*, and *Bullera*. These isolates fell into two groups with identical ITS-LSU sequences: group 1 (GenBank accession OQ388269, here described as *T. erigeronia*) and group 2 (OQ388277 and OQ388309, *T. cardamina*). A fourth isolate was obtained from a separate survey conducted in Ross Township, Michigan, USA in July 2021, during which isolate NB124-2 was isolated as an endophyte of hybrid *Sorghum* roots (OQ434130). Additional attempts to isolate related strains were not successful so far, precluding measurement of their abundance in the community. BLASTn searches of the NCBI GenBank Nucleotide Collection using each ITS-LSU sequence identified an isolate from *Prunus persica* flowers which was closely related: CGMCC 2.6218 (GenBank PV981755/PX225953/GCA_054471015.1), which has 100% sequence identity across ITS-LSU, and 99.21% for RPB1, 95.81% for RPB2, 98.08% for CYTB, and 98.78% for TEF1, and 97.21% average nucleotide identity across the genome to NB124-2. This isolate was recently described as *Fanglaniella lipolytica* (Liao et al. 2026). Subsequent whole-genome sequencing, phylogenetic analyses, and physiological tests were performed on the initial representative isolate of each ITS-LSU group. The strains used for these analyses are in Table 1.

**Table 1.** Strains studied.

| Species | Strain | Date | Location | Source | Genome accession |
|---|---|---|---|---|---|
| *Thailandicola erigeronia* | JL201<sup>T</sup> = NRRL 64467 = CBS 18377 | August 27, 2021 | Durham , North Carolina, USA | Leaf of *Erigeron annuus* | GCA_033819885.2 |
| *Thailandicola cardamina* | JL221<sup>T</sup> = NRRL 64468 = CBS 18378 | October 27, 2021 | Durham , North Carolina, USA | Leaf of *Cardamine hirsuta* | GCA_033819875.2 |
| *Thailandicola cardamina* | JL257 = NRRL 64927 = CBS 19403 | January 25, 2022 | Durham , North Carolina, USA | Leaf of *Cardamine hirsuta* |  |
| *Thailandicola lipolytica* | NB124-2 = NRRL 64928 = CBS 19404 | July 24, 2021 | Ross Township, Michigan, USA | Roots of hybrid *Sorghum* | GCA_052696405.1 |

### Whole-genome sequencing and phylogenetic analyses support the recognition of two novel species

Genome assemblies were generated for strains JL201, JL221, and NB124-2 by combining Oxford Nanopore and Illumina sequencing (Table 2). The final assemblies of JL201, JL221, and NB124-2 are 17.591 Mbp, 19.439 Mbp, and 18.061 Mbp in length with N50 values of approximately 1.1-1.3 Mbp. Many contigs contained telomeric repeats at one or both ends (<u>Table S3</u>, <u>Figure S1</u>). BUSCO assessment of genome completeness using the Microbotryomycetes odb12 dataset recovered 92.5, 91.9, and 92.7% complete BUSCO genes in JL201, JL221, and NB124-2 assemblies, respectively. Analysis of GC content revealed differences across genomes, with JL221 showing a lower GC content (at 49.56%) than JL201 and NB124-2 (54.56% and 55.70%, respectively). Pairwise average nucleotide identity between the genomes as determined by OrthoANI (Lee et al. 2016) ranged from 69.13% to 72.44% (<u>Table S4</u>) indicating substantial genomic divergence among the three taxa and supporting their separation as distinct species (Lachance et al. 2020).

**Table 2.**
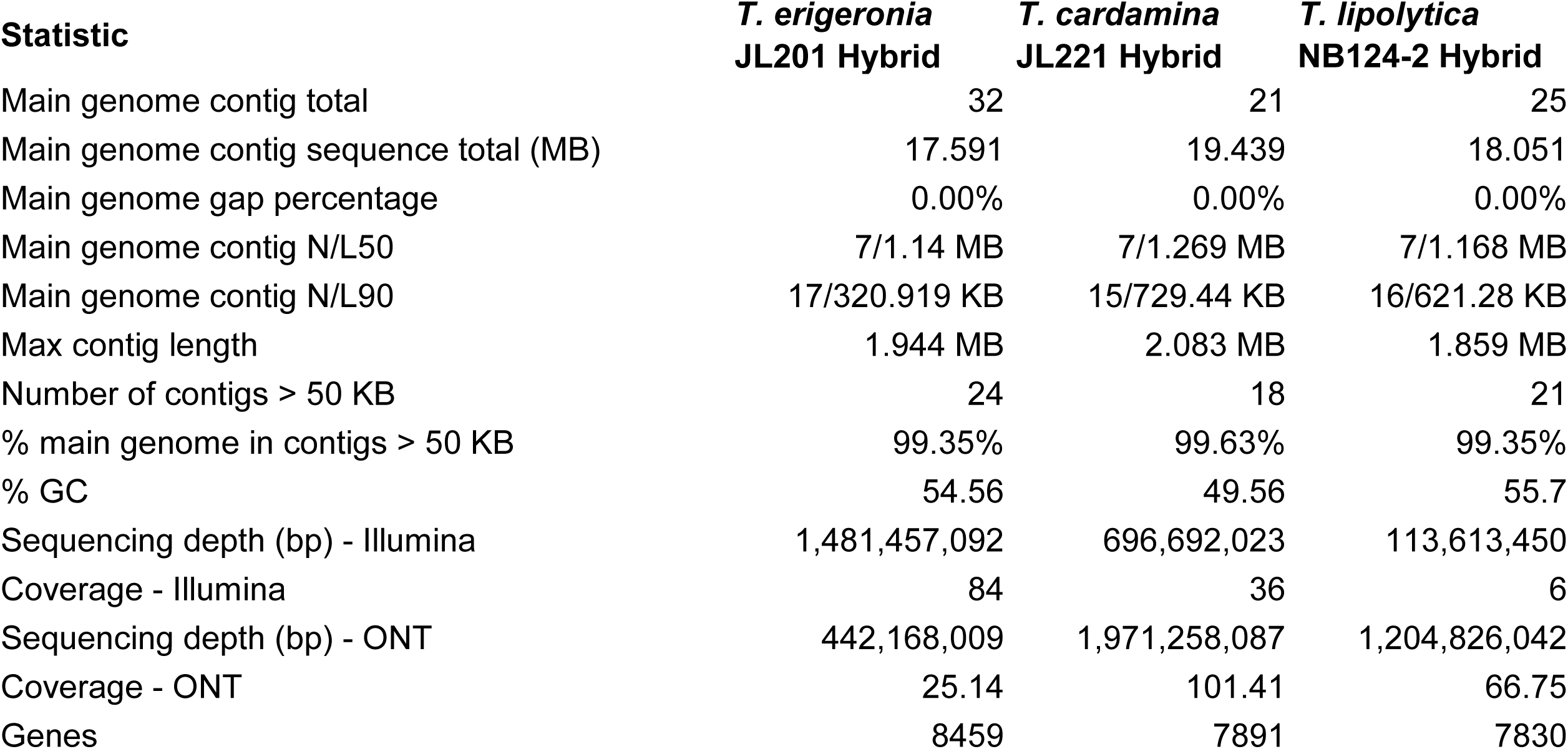
Genome sequencing and assembly statistics.

### Phylogenetics

During the assembly of the phylogenetic dataset for the Microbotryomycetes, several publically available loci sequences were identified as likely contaminants because their best BLASTn matches corresponded to distantly related fungi, and were thus excluded. Additionally, one taxon, *Reniforma strues* (Pore and Sorenson 1990) was examined in greater detail because it showed both an unusually long branch in preliminary analyses and inconsistent placement of its sequenced loci when compared against type-strain sequences. Jiang et al. (2024) also observed long branch length and inconsistent placement of *R. strues*. To determine whether the publicly available sequences were all from the same organism and to verify its phylogenetic placement, we sequenced and assembled the genome of the type strain (NRRL Y-17275). The genome-derived sequences matched those previously deposited in GenBank. We then included the assembled genome in our genome-scale coalescent tree inference and later seven-locus phylogeny inference, which placed *R. strues* within a clade containing the Atractiellales, Heterogastridiales, and *Trigonosporomyces*.

The seven-locus genome-restrained phylogeny (Figure 1) of Microbotryomycetes places all three isolates in a fully supported monophyletic clade sister to the Microbotryales (including *Microbotryum, Microbotryozyma*, *Bauerozyma*, and *Ustilentyloma*) and *Yuzhangia*. Within this clade are the recently described *Thailandicola limtongiae*, *T. hydei*, and *Fanglaniella lipolytica* (Kodchasee et al. 2026; Liao et al. 2026). The clade containing our isolates, *Thailandicola*, and *Fanglaniella* and the Microbotryales clade have a high confidence placement in a clade with *Sampaiozyma*. Furthermore, these clades are inferred to be sister to *Leucosporidium*. A clade supported at 100% places the Sporiodiobolales (*Rhodotorula*, *Rhodosporidiobolus*, and *Sporobolomyces*) with *Curvibasidium*, *Pseudoleucosporidium*, *Nakaseozyma*, *Baiomyces*, and *Fengyania*. A partially-supported (1/82%) clade places the Sporiodiobolales and sister taxa with the clade containing *Leucosporidium*, *Sampaiozyma*, the Microbotryales, *Yuzhangia*, and the novel isolates.

**Figure 1.**
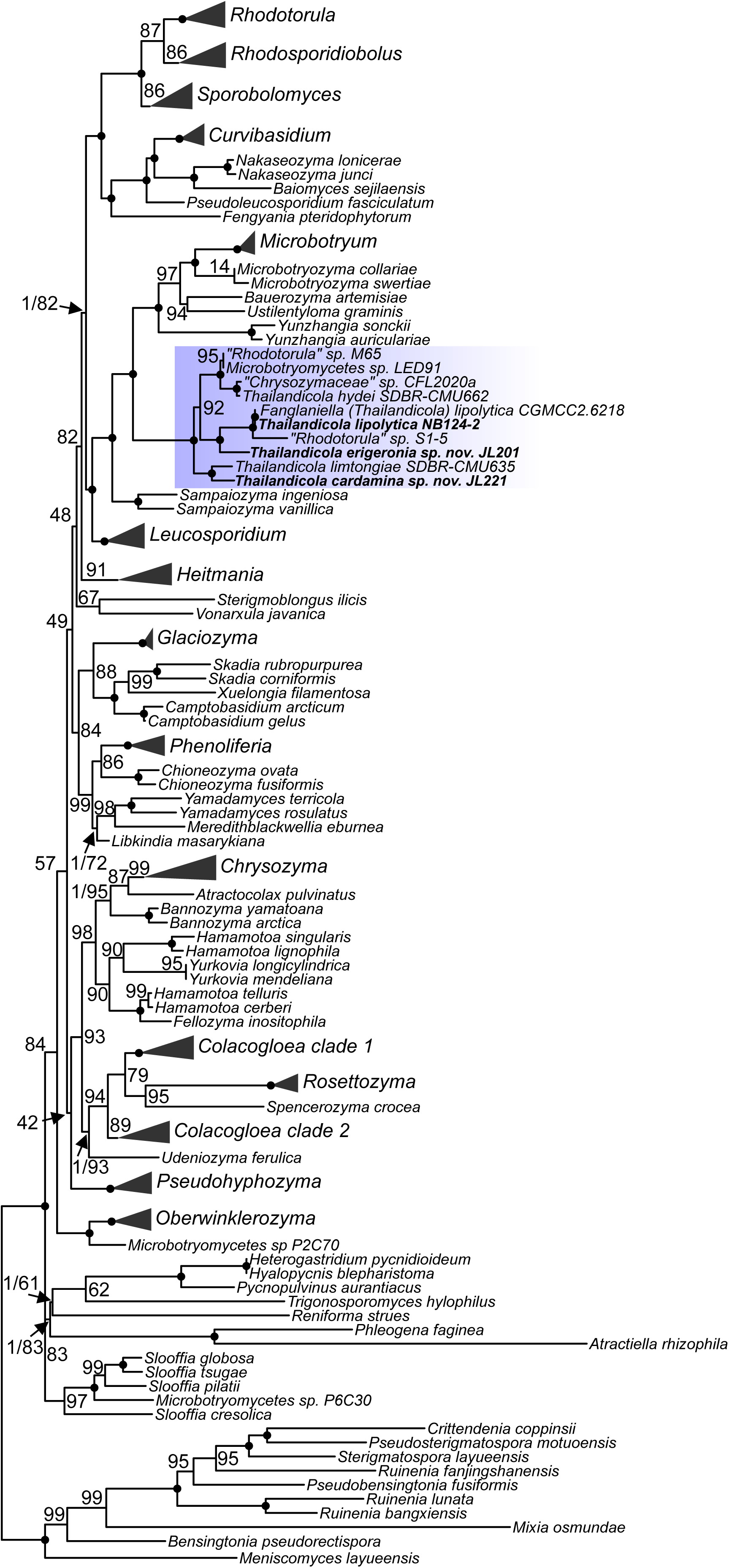
Multi-locus phylogeny of maximum likelihood phylogeny of Microbotryomycetes. Phylogeny of Microbotryomycetes inferred in two steps with an ASTRAL tree generated with whole genomes, then used to constrain a seven-locus (SSU, LSU, 5.8S, *CYTB*, *TEF1*, *RPB1*, and *RPB2*) maximum likelihood phylogeny generated with IQ-TREE. Node support is indicated by solid circles for 100% ultrafast bootstrapping (UFBoot2) support and 1.0 local posterior probability (LPP; if determined in ASTRAL). Otherwise, the UFBoot2 value and LPP are displayed (LPP/UFBoot2 or UFBoot2). Genera which were inferred as monophyletic clades are collapsed for display, and the genus of interest (*Thailandicola*) is highlighted in blue.

Because of the placement of the isolates in a fully-supported monophyletic clade inseparable from the currently described species of *Thailandicola*, we nominate these isolates as two novel species in the genus *Thailandicola*: *T. erigeronia* (JL201) and *T. cardamina* (JL221 and JL257). A third isolate, NB124-2, appears to be conspecific with *Fanglaniella lipolytica* via average nucleotide diversity (ANI, 97.2%) and the placement of both of these isolates suggests that this species should be transferred to *Thailandicola*. As predicted by the average nucleotide diversity, NB124-2 (now *T. lipolytica*) and *T. erigeronia* with 72.4% ANI were inferred to form an exclusive clade, with *T. cardamina* (69.4 and 69.1% ANI to *T. lipolytica* and *T. erigeronia*) more distant and sister to *T. limtongiae*. We also identify 4 isolates with deposited sequences as being placed within *Thailandicola*: “*Rhodoturula*” sp. M65 (JQ956421), Microbotryomycetes sp. LED91 (PZ326142), “Chrysozymaceae” sp. CFL2020a (MW414528), and “*Rhodotorula*” sp. S1-5 (MW534871/MW534894), none of which appear conspecific with described species. The backbone of the Microbotryomycetes is composed of several short branches, many with low confidence, which preclude us from making durable family- and order-level taxonomic placements. However, the Microbotryomycetes are fully supported as a monophyletic clade.

### Comparative Genomics

A comparative genomics approach was used to compare the predicted proteomes of *Thailandicola* spp. with predicted proteomes of other members of Microbotryomycetes (Figure 2), in the genera *Heterogastridium*, *Hyalopycnis*, *Leucosporidium*, *Meredithblackwellia*, *Microbotryum*, *Phenoliferia*, *Rhodotorula*, and *Sporidiobolus*, as well as Microbotryomycetes spp. P2C70 and P6C30 from JGI Mycocosm, which are likely *Oberwinklerozyma* sp. and *Slooffia* sp., respectively, given the inferred phylogeny. These proteomes were functionally annotated using the InterProScan pipeline for both *Thailandicola* and the publically-available datasets. However, genome assemblies and gene model annotations were performed independently and from variable types of sequencing data. In *Thailandicola* spp. the most strongly depleted annotation terms included ribonuclease inhibitors (and RNI-like proteins), proteins containing MYND-type zinc finger domains, furins, F-box-like proteins, and proteins with a proline rich extensin signature (Figure 2A). Conversely, proteins likely originating from transposons were the most enriched in *Thailandicola* spp., including integrases, reverse transcriptases, polyproteins, and DNA/RNA polymerases. These annotations were clustered in at most a single site per contig (<u>Figure S1</u>), suggesting that they may be localized to centromeres. When examining differences within the genus (Figure 2B), this enrichment of transposon components is most pronounced in *T. cardamina* JL221 (69 integrase and 108 reverse transcriptase Pfam domains), notably depleted in *T. lipolytica* NB124-2 (8 and 55), and slightly depleted in *T. erigeronia* JL201 (27 and 51). The most notably metabolic annotations which differ are those for ATP citrate synthase/citrate lyase, which shows some enrichment in *T. cardamina* JL221.

**Figure 2.**
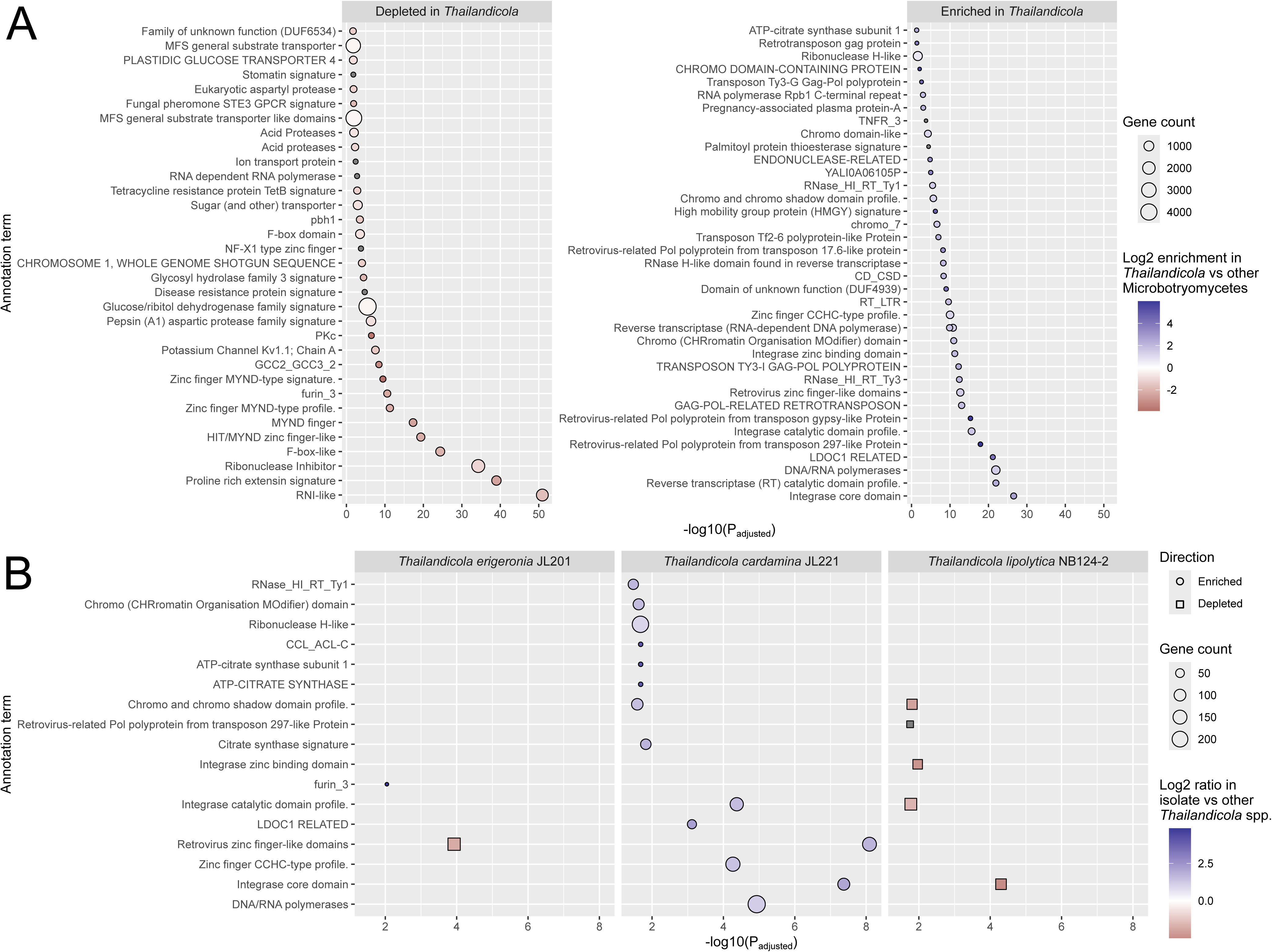
Enrichment of annotation terms in *Thailandicola* and its species. Annotation terms derived from InterProScan were compared in relative proportion between *Thailandicola* and twelve other members of the Microbotryomycetes (A) or between each species and the other two within *Thailandicola* (B). Significance was determined with Fisher’s exact test, and P values were adjusted within each analysis.

### Yeast Phenotypes

Yeast cultures streaked onto plates appeared generally similar, as cream colored, entire margin, and butyrous (Figure 3). Phenotypic characterization of the novel yeast species *T. erigeronia* and *T. cardamina* revealed that these yeasts share a very similar macro- and micromorphology, absence of hyphae or pseudohyphae, absence of fermentation of any tested carbon sources, generally similar carbon and nitrogen assimilation profiles, and similar tolerance to acetic acid, cycloheximide, and osmolytes, and share tolerated ranges of pH and temperature. However, notable differences in each species’ vitamin requirements and the ability to assimilate some specific carbon sources are observed (D-mannitol, L-malic acid, D-ribose, L-rhamnose, and Tween 80). The full set of analyzed traits is found in <u>Table S5</u>.

**Figure 3.**
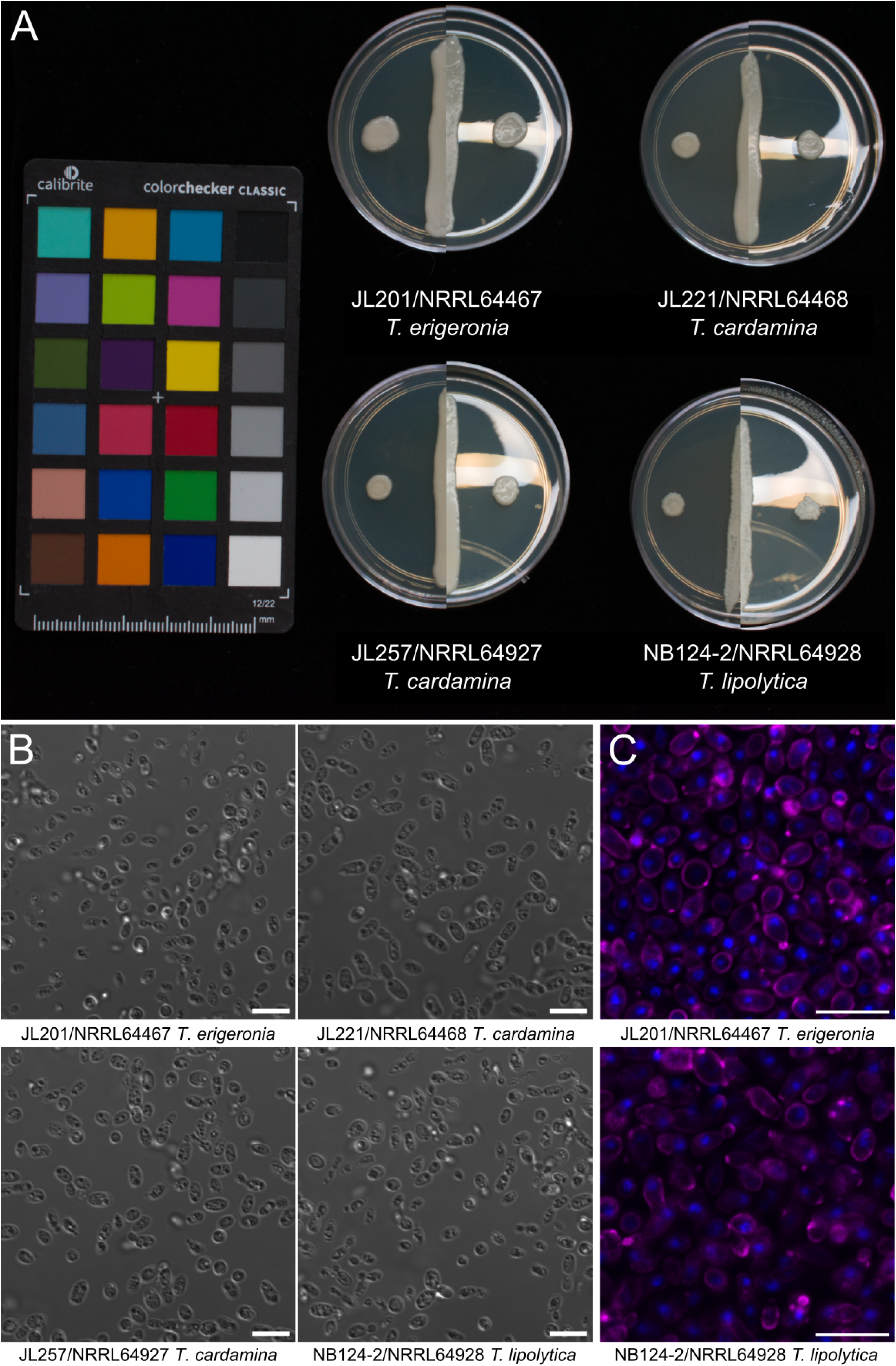
Macro- and micromorphology of *Thailandicola* strains. (A) Composite of colony macromorphology after 4 days of incubation on YMA at 25°C, using both reflecting and non-reflecting lighting. (B) DIC micrograph demonstrating cell micromorphology of planktonic cells grown in YMB for 5 days at 25°C. (C) Cells stained with the DNA stain DAPI (blue) and the cell-wall binding stain wheat germ agglutinin (magenta; WGA-640R) observed with confocal laser scanning microscopy. (B,C) Scale bars measure 10 μm.

### Genetic Transformation

Genetic transformation of two isolates of *Thailandicola* spp. is demonstrated by PCR and heterologous expression of eYFP (Figure 4). Each transformation produced 100s to 1000s of colony forming units, of which a minority showed strong expression of the fluorescent marker. Mapping of the insertion locations with splinkerette PCR was performed for first the right border of the T-DNA construct, then the sites were attempted to be confirmed by PCR amplification across the insertion locus. While this mapping was successful for *T. erigeronia* JL201, the absence of any band for *T. lipolytica* NB124-2 led us to sequence from the left border using ONT linear amplicon sequencing. Of 2,931 reads, a single long read mapped outside the left border into contig 3 at position 527,250 (total contig length of 1,329,339 bp), a different large contig from the right border, which mapped to position 190,852 on contig 20 (total contig length of 329,703 bp). The mapping of reads to the middle of two large contigs suggest that a chromosome translocation occurred during the transformation, and the large number of reads extending into the plasmid sequence after the left border suggests that multiple copies of concatenated T-DNA were inserted into the genome.

**Figure 4.**
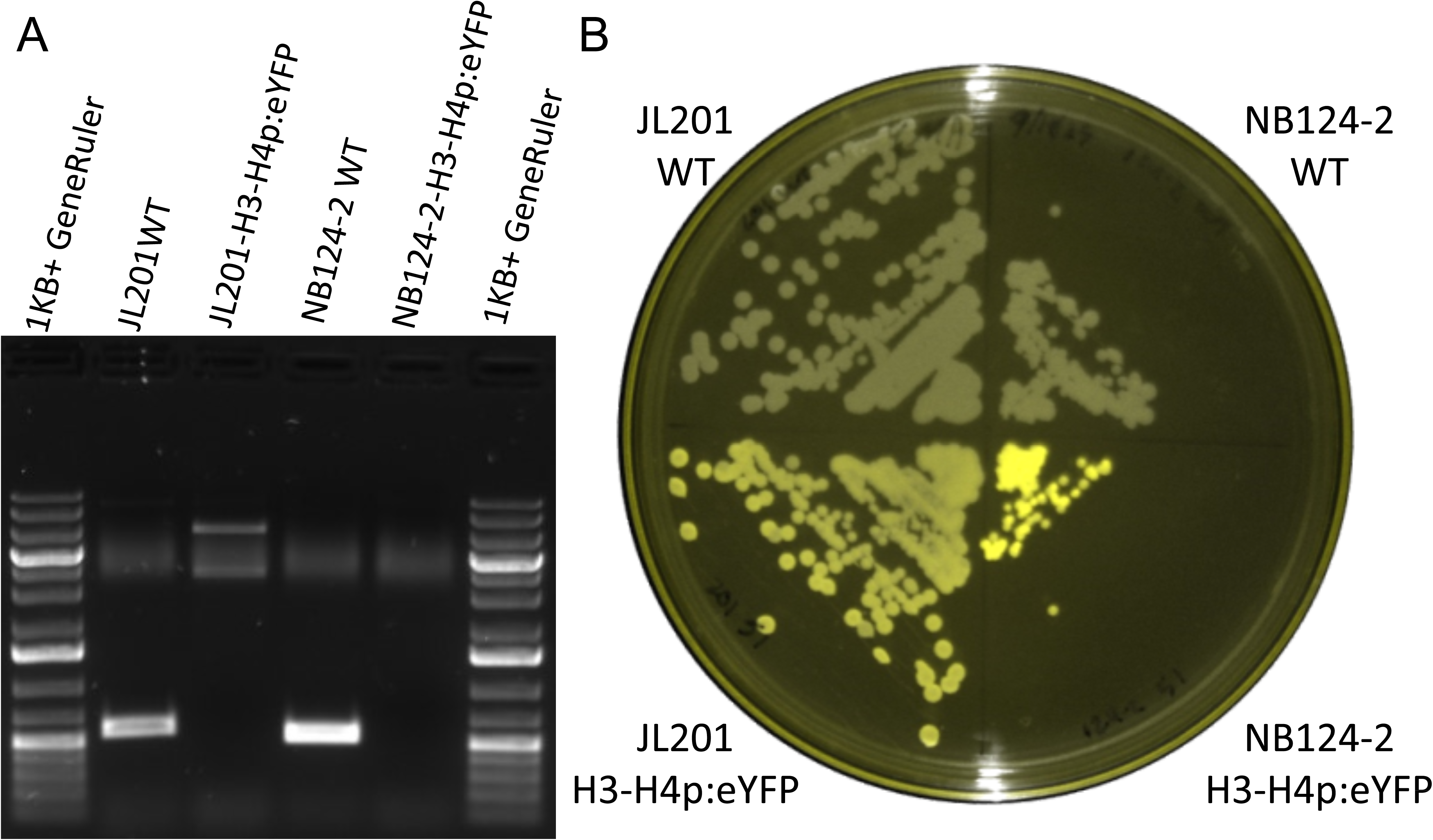
Transformation of strains JL201 and NB124-2. Isolates JL201 (*T. erigeronia*) and NB124-2 (*T. lipolytica*) were genetically transformed to express eYFP via *Agrobacterium tumefaciens*. (A) PCR using primers bridging mapped insertion sites for wild-type and transformed lines. The double bands in lane 3 in the JL201 transformant may be due to concatenated T-DNA insertions in this line, and possible induction of a local aneuploidy. The absence of the band in lane 5 in the NB124-2 transformant appears to be due to both concatenated T-DNA insertions at the locus and likely chromosomal translocation, evidenced by Sanger sequencing from the left and right borders. (B) False-color fluorescence signal of transformed colonies, captured using an iBright™ FL1500 Imaging System with excitation filter range 455-485nm and emission range 508-537nm.

### Mating Locus

Mating compatibility in basidiomycetes is often governed by two *MAT* loci, the pheromone/receptor (*P/R*) locus and the homeodomain (*HD*) locus. When these loci are unlinked, the system is tetrapolar, whereas bipolar systems result when they become linked or when one locus no longer determines mating type (Fraser et al. 2007). For the three examined species, only *T. lipolytica* has the *HD* and *P/R* loci on separate telomere-to-telomere contigs, supporting a tetrapolar mating system in which each locus segregates independently during meiosis (Figure 5). However, the near-chromosome-scale contigs of *T. cardamina* and *T. erigeronia*, together with their high degree of synteny with *T. lipolytica*, also support the inference of a tetrapolar system in these two additional species. Notably, *T. lipolytica* NB124-2 and CGMCC 2.6218 have over 95% identity across their *HD* and *P/R* loci and thus we will discuss based on the more contiguous NB124-2 assembly. Furthermore, the pheromone precursor gene was not assembled in the CGMCC 2.6218 assembly.

**Figure 5.**
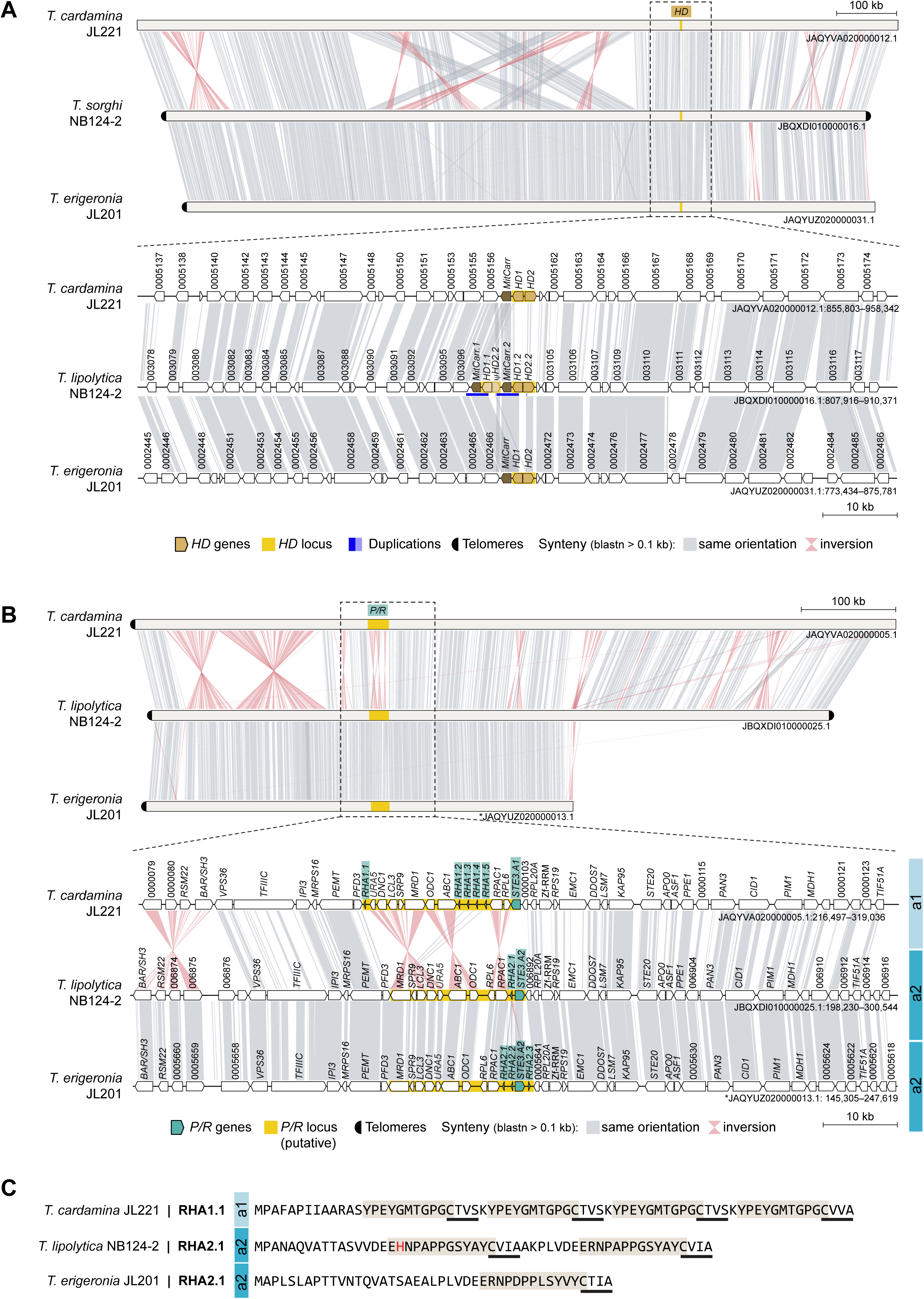
Organization and comparison of the *MAT* loci in three *Thailandicola* species. (A,B) Comparison of the *HD* locus (A) and the putative *P/R* locus (B) in three *Thailandicola* species. In each panel, the top subpanel shows a synteny view of the full contig, highlighting the chromosomal location of the focal locus, and the bottom subpanel provides a zoomed-in synteny view centered on that region. Conserved syntenic blocks are connected by shaded ribbons, with gray indicating the same orientation and pink indicating inversions. Locus-specific genes and the corresponding locus are highlighted as indicated in the key, together with other annotated features. In panel A, the *HD* region of *T. lipolytica* contains two partially identical *HD1* copies that differ at their 5′ ends, consistent with a local duplication event (blue-highlighted region), and a partial *HD2* fragment fused to a neighboring flanking gene. Chromosomes inverted relative to their original assembly orientations are marked with asterisks. (B) The P/R locus of *T. lipolytica* and *T. erigeronia* maintain substantial synteny within the locus and sequence conservation of *STE3*, suggesting that they originate from the same *a2* allele, while *T. cardamina* has multiple regions of inverted similarity and low nucleotide sequence conservation of *STE3* relative to *T. lipolytica* suggesting origin from a different (*a1*) allele. (C) Comparison of pheromone precursor proteins encoded in the *P/R* region. Repeated sequence segments proposed to correspond to the peptide moiety of the mature pheromone are shaded, and sequences resembling the C-terminal CAAX prenylation motif are underlined.

All three species encode an *HD* locus with broadly similar structure, composed of the typical divergently transcribed *HD1* and *HD2* homeodomain genes (Figure 5A). However, the *HD* region of *T. lipolytica* includes two partially identical *HD1* copies that differ at their 5’ ends, consistent with a local duplication event, as well as a partial *HD2* fragment fused to a neighboring flanking gene. Aside from this feature, the *HD* locus sequences are more similar between *T. lipolytica* and *T. erigeronia* than between either of those species and *T. cardamina*, consistent with their closer relatedness.

A similar pattern is also apparent at the *P/R* locus, where *T. lipolytica* and *T. erigeronia* carry the *P/R a2* allele, whereas *T. cardamina* carries the opposite *P/R a1* allele, as inferred from sequence similarity of both the predicted receptor and mating pheromone (Coelho et al. 2008; Maia et al. 2015; Xu et al. 2016) (Figure 5B). The inferred *a1* and *a2 P/R* alleles also differ in gene order and orientation across the compared regions, with several inversions apparent in the synteny plots. Because these comparisons were performed among different species representing opposite inferred mating types, rather than opposite mating-type strains within each species, the precise boundaries and full gene content of the *P/R* locus in each species should be considered provisional. Despite this caveat, several genes within the predicted *P/R* locus regions of *Thailandicola* spp. were also identified in the *P/R* loci of both Sporidiobolales (Coelho et al. 2010; Liu et al. 2025) and Leucosporidiales (Maia et al. 2015), suggesting a shared evolutionary origin.

Variation within the *P/R* region was also evident among *T. lipolytica* and *T. erigeronia*, which share the same inferred *a2* allele. *T. lipolytica* contains a single pheromone gene (*RHA2*) that potentially encodes two peptide repeats, one differing from the other by a single amino acid, whereas *T. erigeronia* contains three identical pheromone genes, each encoding only a single copy of the predicted peptide moiety. By contrast, *T. cardamina* contains five identical pheromone genes (*RHA1*) at the *P/R* locus, each encoding four copies of the predicted peptide moiety. Despite evidence of a heterothallic mating system, the lack of additional known isolates of these species precludes experimental validation of the mating system.

## Taxonomy

### Thailandicola erigeronia Liber, Coelho, & He, sp. nov

#### MycoBank ###

Etymology: growing in/on *Erigeron annuus*.; erigeronia = of Erigeron (fleabanes)

**Type**: USA. NORTH CAROLINA: Durham County, Durham, Duke University West Campus, 36.00°N, 78.93°W, 100 m elevation, ballistospore isolate from *Erigeron annuus* leaf in vegetated strip along road, August 27, 2021. J.A. Liber strain JL201 (**Holotype** NRRL 64467, preserved in a metabolically inactive state, **isotype** CBS 18377). GenBank: OQ388269 (ITS), GCA_033819885.2 (Genome assembly).

Culture characteristics: Colonies from streak culture are cream colored, butyrous, smooth, and without pseudohyphae on YPD. Cells grown in YM broth for 5 days are ellipsoidal to ovoid, 5.34±0.61 μm long by 4.29±0.47 μm wide (mean ± standard deviation, n=51), with polar budding from a narrow base. No evidence of pseudohyphae in Dalmau culture. Absence of ballistospores after 3 weeks on CMA at 20-22°C.

Physiological and biochemical characteristics: Negative for fermentation of D-glucose, D-galactose, sucrose, lactose, raffinose, and D-xylose. Positive for assimilation of D-glucose, L-sorbose, D-glucosamine, sucrose, maltose, α,α-trehalose, methyl-α-D-glucoside, raffinose, melezitose, starch, glycerol, D-mannitol, 2-keto-D-gluconate, D-gluconate, succinate, citrate, palatinose, L-malate, and Tween 80 as sole carbon sources. Weak or delayed assimilation of D-galactose, D-xylose, D-arabinose, cellobiose, salicin, arbutin, inulin, xylitol, DL-lactate, and gentiobiose as sole carbon sources. D-ribose, L-arabinose, L-rhamnose, melibiose, lactose, erythritol, myo-inositol, methanol, and ethanol are not assimilated. Positive for assimilation of ammonium sulfate as a sole nitrogen source. Weak or delayed assimilation of L-lysine hydrochloride, cadaverine hydrochloride, glucosamine, and D-tryptophan as sole nitrogen sources. Potassium nitrate, sodium nitrite, creatine, creatinine, and imidazole are not assimilated. Growth in vitamin-free medium, but reduced growth in absence of thiamine hydrochloride. Positive for urease activity, negative for extracellular starch formation and production of acetic acid. Negative for growth with 1% acetic acid, 16% NaCl, or 60% (w/v) D-glucose (w/v), or 0.1% cycloheximide. Positive for growth in 10% NaCl, 50% (w/v) D-glucose, or 0.01% cycloheximide. Maximum growth temperature is 30°C. Reduced growth at pH=3. No growth at pH=9. Weak liquefaction of gelatin.

Habitat and distribution: Isolated from *Erigeron annuus* leaves in Durham, NC, USA, and not found in other databases.

Diagnosis: Physiologically, *T. erigeronia* differs from the closely related species *T. cardamina, T. limtongiae* and *T. lipolytica* by its ability to assimilate D-mannitol and L-malic acid. While *T. hydei* is also able to assimilate D-mannitol, *T. erigeronia* is able to assimilate L-sorbose, raffinose, soluble starch, and melezitose which *T. hydei* cannot. Morphologically, its cells are more rounded at a length:width ratio of 1.24.

Notes: This isolate appears to be the only representative of the species known, given current searches of databases.

### Thailandicola cardamina Liber, Coelho, & He, sp. nov

#### MycoBank ###

Etymology: growing in/on *Cardamine hirsuta*; cardamina = of Cardamine (bittercresses) Typification: USA. NORTH CAROLINA: Durham County, Durham, Duke University West

Campus, 36.00°N, 78.94°W, 110 m elevation. Ground whole unsterilized leaf tissue of *Cardamine hirsuta* plated on PDA, from a grassy vegetated strip along a road. October 27, 2021. J.A. Liber strain JL221 (**Holotype** NRRL 64468, preserved in a metabolically inactive state, **isotype** CBS 18378). GenBank: OQ388277 (ITS), GCA_033819875.2 (Genome assembly).

Culture characteristics: Colonies from streak culture are cream colored, butyrous, smooth, and without pseudohyphae on YPD. Cells grown in YM broth for 5 days are oblong to ovoid, 6.13±0.72μm long by 3.52±0.46 μm wide (mean ± standard deviation, n=51), with polar budding from a narrow base. No evidence of pseudohyphae in Dalmau culture. Absence of ballistospores after 3 weeks on CMA at 20-22°C.

Physiological and biochemical characteristics: Negative for fermentation of D-glucose, D-galactose, sucrose, lactose, raffinose, and D-xylose. Positive for assimilation of D-glucose, D-ribose, L-rhamnose, sucrose, α,α-trehalose, raffinose, starch, glycerol, 2-keto-D-gluconate, D-gluconate, succinate, citrate, palatinose as sole carbon sources. Weak or delayed assimilation of D-galactose, L-sorbose, D-glucosamine, D-xylose, L-arabinose, D-arabinose, maltose, methyl-α-D-glucoside, cellobiose, salicin, arbutin (variable), melezitose, inulin, DL-lactate, and gentiobiose as sole carbon sources. Negative for assimilation of melibiose, lactose, erythritol, xylitol, D-mannitol, myo-inositol, methanol, ethanol, L-malate, and Tween 80. Positive for assimilation of ammonium sulfate and glucosamine (variable) as sole nitrogen sources. Weak or delayed assimilation of L-lysine hydrochloride, cadaverine hydrochloride, D-tryptophan. Potassium nitrate, sodium nitrite, creatine, creatinine, and imidazole are not assimilated. Growth in vitamin-free medium is negative, but is restored with addition of thiamine hydrochloride. Positive for urease activity, negative for extracellular starch formation and production of acetic acid. Negative for growth with 1% acetic acid, 16% (w/v) NaCl, or 60% (w/v) D-glucose (w/v), or 0.1% cycloheximide. Positive for growth in 10% (w/v) NaCl, 50% (w/v) D-glucose, or 0.01% cycloheximide (weak). Maximum growth temperature is 30°C. Reduced growth at pH=3. No growth at pH=9. Strong liquefaction of gelatin.

Habitat and distribution: Isolated from *Cardamine hirsuta* leaves in Durham, NC, USA, and not found in other databases.

Additional material examined: strain JL257 (NRRL 64927, CBS 19403) Durham County, Durham, Duke University West Campus, 36.00°N, 78.94°W, 114 m elevation. Ground whole unsterilized leaf tissue of *Cardamine hirsuta* plated on PDA, from a grassy vegetated strip along a road. January 25, 2022. J.A. Liber.

Diagnosis: Physiologically, *T. cardamina* differs from the closely related species *T. erigeronia, T. limtongiae,* and *T. lipolytica* by its ability to assimilate L-rhamnose and its inability to grow in vitamin-free medium. *T. hydei* can also grow on L-rhamnose, but is unable to grow on L-sorbose, D-arabinose, raffinose, or soluble starch on which *T. cardamina* can grow. Morphologically, its cells are more ellipsoidal than either species with a length:width ratio of 1.74.

Notes: These two isolates appear to be the only ones known, given current searches of databases.

### *Thailandicola lipolytica* (A.H. Li & Y.J. Liao) Liber, Coelho, & He *comb. nov*

#### MycoBank MB573041

Basionym: *Fanglaniella lipolytica* A.H. Li & Y.J. Liao, Journal of Fungi 12: 7 (2026)

## Discussion

Here, we present the description of a novel basidiomycete yeast genus containing two novel species from three isolates and a closely related third species, *Thailandicola lipolytica*, all isolated from plant leaves and roots. These isolates are characterized for their morphology, metabolism, stress tolerance, genomic sequence and overlying gene models, and inferred phylogenetic relationships. The combination of these lines of evidence supports the description of two novel species within the recently-described genus *Thailandicola* in the Microbotryomycetes.

The inferred phylogeny generally shows a similar structure to that inferred by Jiang et al. (2024), but some notable differences were observed. Several nodes that were poorly supported in previous studies (Wang et al. 2015; Li et al. 2020; Jiang et al. 2024) are more strongly resolved here. One of the clearest examples is the Microbotryales, which itself is well supported and also forms a well-supported clade with *Sampaiozyma* and *Thailandicola*. Microbotryales was originally described to contain *Microbotryum* and *Ustilentyloma* (Bauer et al. 1997), which is now supported by both multi-locus and genomic data. Earlier topologies in Wang et al. (2015) separated *Sampaiozyma* and *Leucosporidium* from the Microbotryales, while *Heterogastridium* and *Reniforma* were placed as sister taxa with low branch support. Li et al. (2020) and Jiang et al. (2024) inferred a topology similar to the present grouping, albeit with low support, and still including *Reniforma* as sister to *Yuzhangia*. The Sporidiobolales (*Sporobolomyces, Rhodotorula, and Rhodosporidiobolus*) is placed in our phylogeny with *Curvibasidium*, *Pseudoleucosporium*, *Nakaseozyma*, *Baiomyces*, and *Fengyania* with a high degree of support, which may also suggest either re-circumscription of Sporidiobolales or the need for additional family- and/or order-level taxonomic description for these genera.

The improved support for the Microbotryales clade and the nearby placement of *Leucosporidium* and *Sampaiozyma* may partly reflect additional data informing the placement *Reniforma strues*. It is possible that the long branches on which *Yuzhangia* is placed attracted the highly diverged *R. strues* to this clade in earlier analyses (Wang et al. 2015; Li et al. 2020; Jiang et al. 2024). Through sequencing the genome of *R. strues*, we placed this species in a clade distant from the Microbotryales. This clade contains *Trigonosporomyces*, the Atractiellales, and the Heterogastridiales. *Trigonosporomyces* and Heterogastridiales are currently placed within Microbotryomycetes, while the Atractiellales are within their own class (Atractiellomycetes). Given the inferred phylogeny, the transfer of the Atractiellomycetes to Microbotryomycetes would be justified.

Well-supported nodes suggest the placement of the novel species and *Fanglaniella* within *Thailandicola* which forms a clade with the Microbotryales and *Sampaiozyma*. In the recent work erecting *Thailandicola,* the authors used multi-locus data from the same loci that were used for our phylogenetic inference (Kodchasee et al. 2026), but did not include genomic-scale data. Our approach to include loci across the genome with a genome-constraint tree again found support for *Thailandicola* as a monophyletic clade, further solidifying this genus. Morphologically, descriptions of *Fanglaniella* (Liao et al. 2026) and *Thailandicola* (Kodchasee et al. 2026) are consistent to those we describe here, with polar budding, ellipsoidal to ovate cells, and cream to pale yellow streaks. However, we did not observe hyphae or pseudohyphae in *T. erigeronia* or *T. cardamina*, while these were observed for *T. hydei* and *T. limtongiae.* In considering the expanded genus, *Thailandicola* is not clearly morphologically distinct from the closely related yeasts such as *Sampaiozyma* and *Yuzhangia*, yet remains distinct from other members of a more inclusive genus that would include plant pathogens with known teleomorphs including *Microbotryum* and *Ustilentyloma* (Wang et al. 2015).Given the phylogenetic and phenotypic evidence, we believe that *Thailandicola* continues to be justified as a reasonable genus for grouping these five species.Furthermore, we continue to find that the Microbotryomycetes are well supported as a class, but the nodes along the backbone show short branch lengths and low support, so that many *incertae sedis* classifications remain for family- and order-level clades.

The relatively low ANI between isolates (∼70%), and distinct metabolic and slight morphological traits, suggest that these isolates represent unique species under the phylogenetic and morphological species concepts. Attempts to induce mating among the isolates were unsuccessful so far, but fungal sexual cycles can be difficult to recapitulate *in vitro*, even among isolates of the same species (Fraser et al. 2003; O’Gorman et al. 2009).

Given our analysis of the mating loci, these isolates appear to be heterothallic, with *a1* and *a2* alleles at the *P/R* locus and a comparatively conserved *HD* locus. The greater structural variation observed in the *P/R* region, including differences in pheromone gene copy number, is consistent with the broader tendency of the *P/R* locus to be more evolutionarily labile than the *HD* locus in basidiomycetes. In this context, it is notable that recent work in *Cryptococcus* linked pheromone gene duplication to structural instability and expansion of the *P/R* region, likely through the generation of inversion-prone architectures (Coelho et al. 2025). In *Thailandicola*, variation in pheromone gene number and organization may likewise reflect a dynamic history of local expansion and contraction within the *P/R* locus. This possibility may also be relevant to other Microbotryomycetes, such as *Leucosporidium scottii* (Maia et al. 2015) and Sporidiobolales (Sen et al. 2019; Liu et al. 2025), which exhibit a broadly similar locus organization.

The comparative genomics approach to examine genome annotations appears to reflect an increased abundance of retrotransposons in *Thailandicola*, given the enrichment of integrase, reverse transcriptase, ribonuclease H, chromo domain, and polyprotein elements (Muszewska et al. 2011) in this genus. However, an important caveat is that the detection of certain functional annotations may be affected by differences in genome sequencing strategy, assembly quality, and gene model prediction across the panel of compared genomes and their respective proteomes. This may be especially relevant for repetitive elements, such as retrotransposon-derived sequences, which are more challenging to detect in fragmented short-read assemblies (Zhou et al. 2020) we cannot exclude the possibility that the lower fragmentation of the *Thailandicola* genomes relative to other available assemblies contributed to the more frequent annotation of such elements. However, many of the other significantly enriched or depleted annotations are not similarly repetitive and thus are less likely to be biased by the quality of the predicted proteome.

Beyond these retrotransposon-associated features, the proteomes of *Thailandicola* spp. are depleted of ribonuclease inhibitors (RIs or RNIs) and RNI-like proteins. While these are not known in the literature to be associated with retrotransposon defense, their anti-ribonuclease activities (Kobe and Deisenhofer 1996) could suggest an interaction with retroviral ribonucleases. Proline-rich extensins, also depleted in *Thailandicola*, are proteins typically known from plant cell walls (Marcus et al. 1991; Saha et al. 2013). However, this same protein family (PR01217) is present in fungal proteins such as the RNA polymerase II RPO21 in *Saccharomyces cerevisiae* (Blum et al. 2025). Due to the presence of this annotated family in proteins of other known functions, the importance of its differential abundance is uncertain. This similarly applies to proteins containing F-box-like and MYND finger domains. ATP-citrate synthase/lyase was notably the only energy metabolism pathway annotation enriched in one of the *Thailandicola* species. This enzyme is involved in lipid production (Chypre et al. 2012), and in the oleaginous yeast *Yarrowia lipolytica* its heterologous expression results in enhanced lipid accumulation (Zhang et al. 2014). While we have not examined the macromolecular composition of these yeast isolates, the relative enrichment of this annotation suggests that *T. cardamina* may favor lipid accumulation compared to the other species.

Since Ruinen (1961) described the phyllosphere as “an ecologically neglected milieu” sixty-five years ago, the surfaces of plants, especially leaves, have been intensely studied with tens of thousands of metagenomic samples collected and deposited into GenBank (Sohrabi et al. 2023). Despite this breadth of sampling, *Thailandicola* spp. are not represented in sequence databases including GenBank and GlobalFungi (Větrovský et al. 2020) outside of the isolates presented here. However this appears not to be unusual for the Microbotryomycetes, for which many species and genera are represented by a single isolate (Li et al. 2020; Jiang et al. 2024). Further, the occurrence of all known isolates of *Thailandicola* from flowers (Kodchasee et al. 2026; Liao et al. 2026), leaves, and roots suggests some degree of adaptation to the plant environment. Closely related taxa including *Leucosporidium* and the Sporidiales are capable of mycoparasitism (Schoutteten et al. 2023), a trait that we did not observe here as these yeasts did not form hyphae with colacosomes, but is frequent in the class and may shape the ecology and distribution of these yeasts.

The demonstrated transformability of this genus is relatively unusual in the literature. Of all basidiomycete yeasts, a handful of taxa have published protocols for transformation including the human pathogens in *Cryptococcus* s.s. (Toffaletti et al. 1993) and the closely related yeast-like *Tremella* (Guo et al. 2008), the red yeasts in Sporidiobolales (Takahashi et al. 2014; Pi et al. 2018), and the plant pathogens in Ustilaginaceae including *Ustilago* (Wang et al. 1988) and *Pseudozyma* (Marchand et al. 2007). Given the potential of many basidiomycete yeasts in industrial (Pi et al. 2018) and agricultural (Filonow 1998) uses, as well as their importance as human pathogens (Kim et al. 2013; Liu et al. 2019), additional genetic tools for understanding their biology are extremely important. While heterologous expression is helpful for some genetic investigations, the ability to disrupt or insert sequence at specific loci is usually necessary for testing the function of genes. This would suggest that further development of CRISPR/Cas9 or homologous recombination approaches like those implemented in other Microbotryomycete yeasts are necessary (Takahashi et al. 2014; Jiao et al. 2019; Schultz et al. 2019), and the ability to express a construct heterologously favors a CRISPR/Cas9 approach.

Agricultural productivity poses among the greatest challenges to satisfying the world’s demands for food while preserving a livable planet (Baldos and Hertel 2014). Phyllosphere and rhizosphere microbiota are increasingly studied as an important contributor to plant’s defenses against biotic and abiotic stresses (Bulgarelli et al. 2013; Sohrabi et al. 2023). Our description of these novel yeasts derived from plant surfaces contributes to the conversion of some of the unknown “microbial dark matter” to described organisms with transcriptome-informed annotated genomes, metabolic and physiologic observations, and genetic tools for future experimental study.

## Data availability

Sequencing and assembly data are accessioned under BioProject PRJNA892096. Scripts, phylogenetic trees, and other data are deposited at https://github.com/liberjul/Yeast_genomes

## Acknowledgements

The authors thank Ian Medeiros and Keaton Tremble for assistance with genome annotations and phylogenetic inference. We also thank Gregory Bonito for sharing the isolate NB124-2. We thank the curators at the USDA and CBS culture collections for the deposition of strains. We appreciate the feedback on the manuscript provided by Joseph Heitman, Rytas Vilgalys, and Teun Boekhout. J.A.L. received funding support from Duke University, the Duke Microbiome Center, and the Triangle Center for Evolutionary Medicine. S.Y.H. is an Investigator at Howard Hughes Medical Institute. This study was also supported by the National Institute of Allergy and Infectious Diseases of the National Institutes of Health under award R01 AI050113-20 (M.A.C.).

## Figure and Table Legends

Table S1. Genomes used for comparative genomics analysis.

Table S2. Accessions of data used in phylogenetic inference.

Table S3. Contig sizes and telomeric repeats.

<u>Table S4. Average nucleotide identity between *Thailandicola* spp. genomes.</u> ANI values are calculated with OrthoANI between the query genome (row) and the subject (column).

<u>Table S5. Metabolic and stress tolerance traits.</u> Test numbers come from the deposit form for the Westerdijk Institute’s CBS collection. For assimilation tests, “+” is defined as >50% the optical density of the D-glucose positive control, while “w” is defined as greater than 20% and less than or equal to 50% of the positive control. Other test criteria are defined according to Suh et al. (2008).

